# Cellular responses to FGF1 are modulated by palmitoylation of the docking protein FRS2α

**DOI:** 10.64898/2025.12.11.693497

**Authors:** Seong An, Yoshihisa Suzuki, Jyotidarsini Mohanty, Francisco Tome, Irit Lax, Joseph Schlessinger

## Abstract

An important mechanism by which receptor tyrosine kinases (RTKs) mediate cellular responses involves the formation of signaling complexes through direct interactions with membrane-associated docking proteins, followed by phosphorylation of multiple tyrosine residues. These docking proteins recruit and activate downstream signaling molecules and enzymes following ligand stimulation. The docking protein FRS2α has been established as a major signaling hub activated by fibroblast growth factors (FGFs), neurotrophic factors, and other extracellular cues. Here, we show that palmitoylation of FRS2α at two sites is essential for stabilizing its myristoylation-dependent association with the plasma membrane. FGF1-induced MAPK activation and other cellular responses are partially restored in cells expressing FRS2α mutants deficient in either one of the two palmitoylation sites. However full restoration of signal strength including MAPK response and other FGF1-induced cellular activities requires palmitoylation at both FRS2α sites. In addition to enhancing signaling robustness, anchoring of FRS2α to the plasma membrane creates a structural platform for assembling multi-protein complexes essential for cytoskeletal reorganization associated with membrane ruffling, macropinocytosis, and other FGF1-induced processes. Finally, we demonstrate that while PC12 cells lacking FRS2α or deficient in FRS2α palmitoylation can proliferate, FGF1-induced neuronal differentiation strictly depends on the palmitoylation of the docking protein.

## Introduction

One of the hallmarks of the mechanism of action of receptor tyrosine kinases (RTKs) is their ability to function as both ligand-stimulated enzymes and as platforms for direct interactions and recruitment of cellular target proteins. RTKs also interact with specific cell membrane-associated docking proteins containing multiple tyrosine phosphorylation sites, forming physical complexes with additional signaling molecules critical for activating cellular signaling pathways by tyrosine phosphorylation and by allosteric mechanisms. For instance, docking proteins IRS1 and IRS2 play pivotal roles in mediating signaling pathways activated by the insulin receptor (IR) and IGF1 receptor (IGF1R)^1^. Likewise, the docking protein FRS2α is central to cell signaling induced by members of the FGF family^2^, as well as by neurotrophic factors such as NGF, BDNF, and GDNF families^3^.

FRS2α, and its related protein FRS2β, have similar domain structures, with an N-terminal myristoylation sequence for membrane association, followed by a PTB (phosphotyrosine binding) domain for receptor interaction and multiple C-terminal tyrosine phosphorylation sites^2^. While FRS2α binds constitutively to FGF receptors (FGFR) at the juxtamembrane region of the receptor, it binds in a ligand-dependent manner to neurotrophic factor receptors (TrkA, TrkB, RET) through a canonical tyrosine phosphorylated NPXYp sequence^4,5,6,7^. Ligand-induced phosphorylation of six tyrosine phosphorylation sites in FRS2α serve to recruit SH2 domains of the adapter protein^2^ Grb2 and the tyrosine phosphatase Shp2, which itself recruits Grb2 when tyrosine phosphorylated^8^. Grb2 subsequently recruits Sos^2^, the guanine nucleotide exchange factor for Ras, as well as the docking protein^9^ Gab1, which recruits phosphoinositide 3-kinase (PI3K), to activate multiple signaling pathways. Notably, FRS2α is also subject to a negative feedback mechanism mediated by MAPK-dependent phosphorylation of eight threonine residues which reduce tyrosine phosphorylation of FRS2α; heterologous MAPK stimulation by other extracellular cues, such as insulin, EGF and PDGF, can contribute to this feedback mechanism^10^.

FRS2α is also essential during embryogenesis. FRS2α−/− mice die by embryonic day 8 (E8) due to defects in anterior-posterior axis formation^11^. Furthermore, FRS2α−/− embryos show loss of function of trophoblast stem cells in the extraembryonic ectoderm that give rise to placental tissues, as well as reduced MAPK activation in this embryonic region at E6 of embryonic development^11^. Mouse embryonic fibroblasts (MEFs) isolated from FRS2α−/− embryos survive and proliferate likely via compensatory mechanisms that bypass the central role played by FRS2α in the control of signaling pathways normally activated by FGF stimulation^12^. There is no obvious phenotype of FRS2α−/− mice^13^, which likely reflects the different tissue and temporal expression patterns of FRS2α and FRS2α^14,15^, since FRS2β can rescue FGF-stimulated MAPK activation in FRS2α−/− MEFs^15^. Supporting the importance of FGFR1–FRS2α signaling, mice with a deletion of the juxtamembrane region or mutations (L423A and V429A) in FGFR1, which are required for binding to FRS2α, die at late embryogenesis (E15.5) or at birth, exhibiting multiple developmental defects, respectively^16,17^. This also confirms that in the absence of FRS2α, alternative cellular signaling pathways can compensate for some of FRS2α loss.

A more recent study showed that in addition to myristoylation, FRS2α is palmitoylated at cysteines 4 and 5 in the N-terminal region, in a manner dependent on N-myristoylation^18^. Palmitoylation involves the attachment of palmitate to cysteine residues via a thioester linkage, which is a reversible process mediated by acylation and deacylation enzymes^19^. Mutation of these two cysteines (C4,5S) or the myristoylation site (G2A) disrupts the membrane localization of FRS2α. The G2A mutant was previously shown to be defective in FGF1 signaling^2,20^. Because cysteine palmitoylation is reversible and membrane association of FRS2α is critical for mediating intracellular signaling, it is possible that cellular responses to FGFs, NGF, and other neurotrophic factors are dynamically regulated by FRS2α palmitoylation.

Here, we begin to explore the role of FRS2α palmitoylation in FGF1-induced cellular responses. FRS2α participates in multiple cellular processes, including cell proliferation, differentiation, cytoskeletal organization, and motility. In this report, we focus specifically on how palmitoylation influences FRS2α membrane association, the recruitment and activation of its cellular targets, and downstream events essential for cell proliferation and differentiation as well as for actin remodeling and macropinocytosis.

## Results

To examine the cellular localization of FRS2α and its mutants, we generated an expression vector encoding FRS2α proteins fused to tdStayGold, a tandem dimer form of the highly photostable and bright fluorescent protein StayGold, which has been shown to label proteins in a monovalent manner^21^.

### Both palmitoylation sites are required for membrane association of FRS2α

The expression vectors for imaging included the coding sequence of tdStayGold connected to the C-terminus of FRS2α and its four possible lipid modification mutants (Fig. 1A): singly palmitoylated (C4S and C5S), palmitoylation deficient (C4,5S), and myristoylation deficient (G2A) FRS2α. These fluorescent FRS2α constructs were transiently expressed in FRS2α−/− MEFs^12^ and imaged by confocal microscopy (Fig. 1B). In confocal optical sections (∼800-nm thick), FRS2α WT could be seen to localize primarily to the plasma membrane, while the FRS2α C4,5S and G2A mutants were cytosolic. To improve visualization of membrane localization of FRS2α WT in flat cells, (Fig.1C, left panel) we used hypotonic solution composed of 4:1 (v/v) diH2O/media that causes mammalian cells of various types to swell and dramatically increase in height without bursting^22^. We found this to be the case with FRS2α−/− MEFs as well, and the increased cell height in hypotonic solution made the membrane localization of FRS2α WT much more obvious (Fig. 1C, right panel).

**Figure 1.**
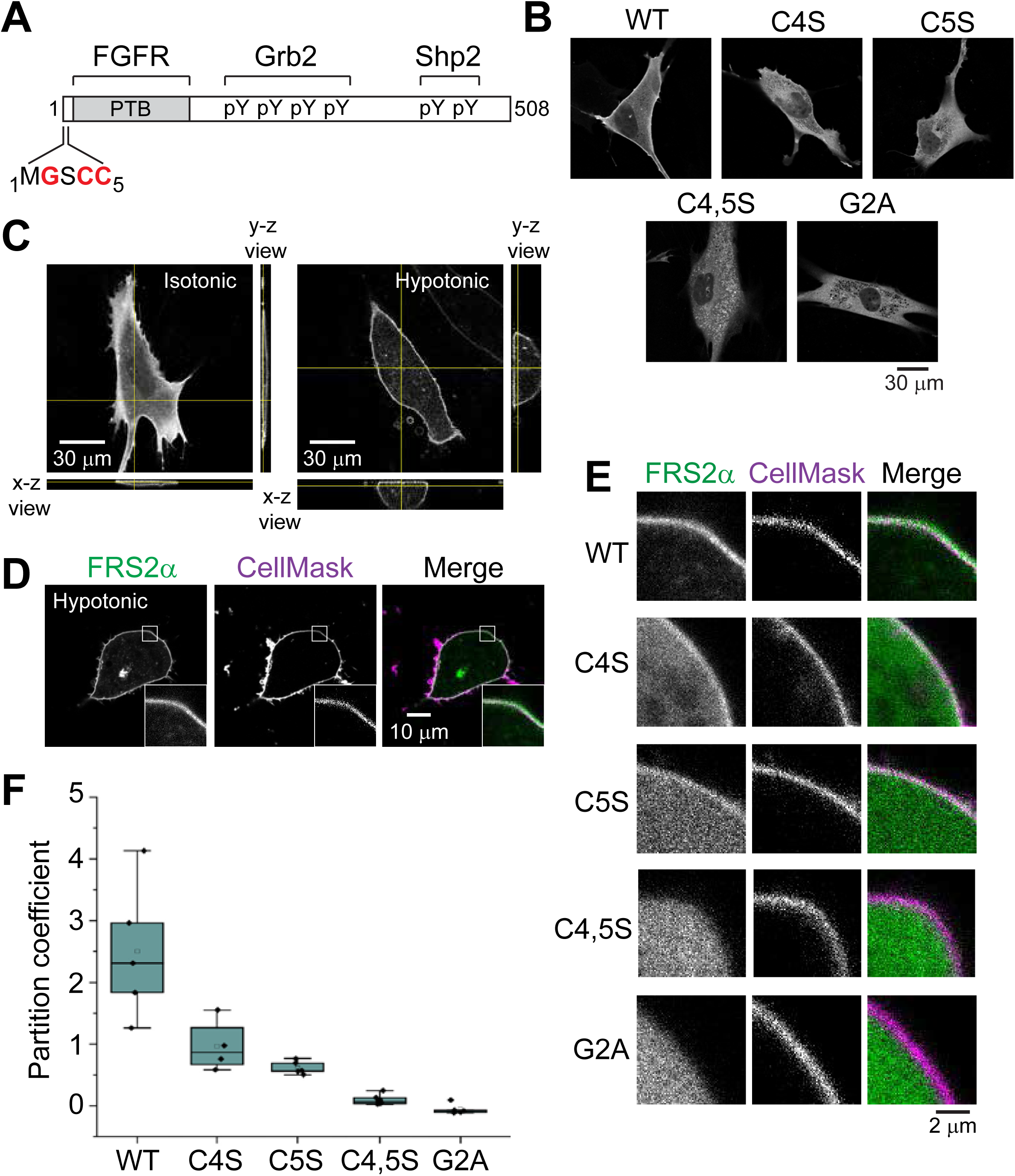
Quantification of membrane binding by FRS2α and its lipid modification mutants. **(A)** Schematic of FRS2α showing N-terminal myristoylation (Gly2) and palmitoylation (Cys4 and Cys5) sites, Phosphotyrosine-binding domain (PTB) that binds to the juxtamembrane region of FGFR, and tyrosine phosphorylation sites that bind to the SH2 domains of Grb2 and Shp2. **(B)** Confocal micrographs of FRS2α−/− MEFs transiently expressing FRS2α WT or its lipid modification mutants fused to tdStayGold at C-terminus. Single confocal planes are shown. **(C)** Orthogonal views (x-z and y-z) of confocal z-stacks of FRS2α WT-tdStayGold in FRS2α−/− MEFs bathed in isotonic or hypotonic media. Yellow lines in x-y views indicate locations of slices for orthogonal projections. Yellow lines orthogonal to the z-axis in side panels indicate confocal planes displayed in x-y views. **(D)** Confocal images of FRS2α WT-tdStayGold in FRS2α−/− MEF stained with the membrane marker CellMask in hypotonic media. Insets show magnified views of boxed regions. **(E)** Magnified views of FRS2α−/− MEFs expressing FRS2α WT, redisplayed from **(D)**, or its lipidation mutants fused to tdStayGold and stained with CellMask in hypotonic media. **(F)** Box plots showing membrane partition coefficients of FRS2α WT or its lipidation mutants fused to tdStayGold in FRS2α−/− MEFs bathed in hypotonic media. The partition coefficient for FRS2α WT is significantly different (P<0.01) from those of C4S and C5S, which in turn are significantly different (P<0.01) from those of C4,5S and G2A.

To quantitatively assess membrane localization of FRS2α WT and its lipid modification mutants, we turned to a method of determining plasma membrane partition coefficients based on modeling linear distribution of membrane and cytoplasm fluorescence across the cell periphery using plasma membrane and cytosolic markers^23^. To this end, we stained cells expressing FRS2α-tdStayGold constructs or tdStayGold alone (Supporting information Fig. 1) with the amphipathic membrane marker CellMask^24,25,26^ and imaged the cells by confocal microscopy in hypotonic solution (Fig. 1D and 1E). In this manner, the membrane localization of FRS2α C4S and C5S could now be seen clearly. Partition coefficients were calculated based on the ratio of plasma membrane and cytosolic fractions. Compared to FRS2α WT, the calculated partition coefficients for FRS2α C4S and C5S were 39% and 25%, respectively (Fig. 1F). For FRS2α C4,5S and G2A, the relative partition coefficients were 4% and −2%, respectively, indicating that while membrane binding is greatly reduced for the C4,5S mutant, it cannot be measured reliably for the G2A mutant, in agreement with the study by Barylko *et al.* using z-scan fluorescence fluctuation spectroscopy^18^. Thus, these experiments demonstrate that full membrane association of FRS2α requires both palmitoylation sites.

It should be noted, as previously reported^18^, that FRS2α C4,5S showed punctate fluorescence within the cell. Based on immunostaining experiments using anti-Rab antibodies, FRS2α C4,5S colocalized to varying degrees with Rab5-positive early, Rab7-positive late and Rab11-positive recycling endosomes (Supplemental Fig. 1); the underlying mechanism of the colocalization is not known.

### Modulation of FGF1 stimulation by palmitoylation of FRS2α

To study the role of FRS2α palmitoylation in FGF1-induced cellular signaling pathways, we stably expressed FRS2α WT or its lipid modification mutants in previously described FRS2α−/− MEFs^12^. We have previously demonstrated that FGF1-induced activation of FGFR or phospholipase-Cγ (PLCγ) does not depend upon FRS2α expression^12^; PLCγ, unlike other FGFR effectors, is directly recruited by the receptor^27^. Accordingly, we first examined whether these early signaling events are affected by the palmitoylation state of FRS2α. Confluent FRS2α−/− MEFs and MEFs rescued with FRS2α WT or its mutants were stimulated with FGF1 or left untreated for 10 min at 37°C. Cell lysates were immunoprecipitated with anti-FGFR1 antibodies, subjected to SDS-PAGE, and analyzed by immunoblotting for tyrosine phosphorylation using anti-pTyr antibodies. Anti-FGFR1 antibodies were used as a loading control (Fig 2A). As expected, we found that none of the lipidation mutations in FRS2α affected tyrosine phosphorylation of FGFR (Fig. 2A, top lane). To examine whether tyrosine phosphorylation of PLCγ is affected, lysates of FGF1-stimulated and untreated cells were immunoprecipitated with anti-PLCγ antibodies and analyzed for tyrosine phosphorylation of PLCγ using anti-pTyr antibodies. Anti-PLCγ antibodies were used as a loading control. As expected, none of the lipidation mutations of FRS2α affected tyrosine phosphorylation of PLCγ (Fig. 2A, second lane).

**Figure 2.**
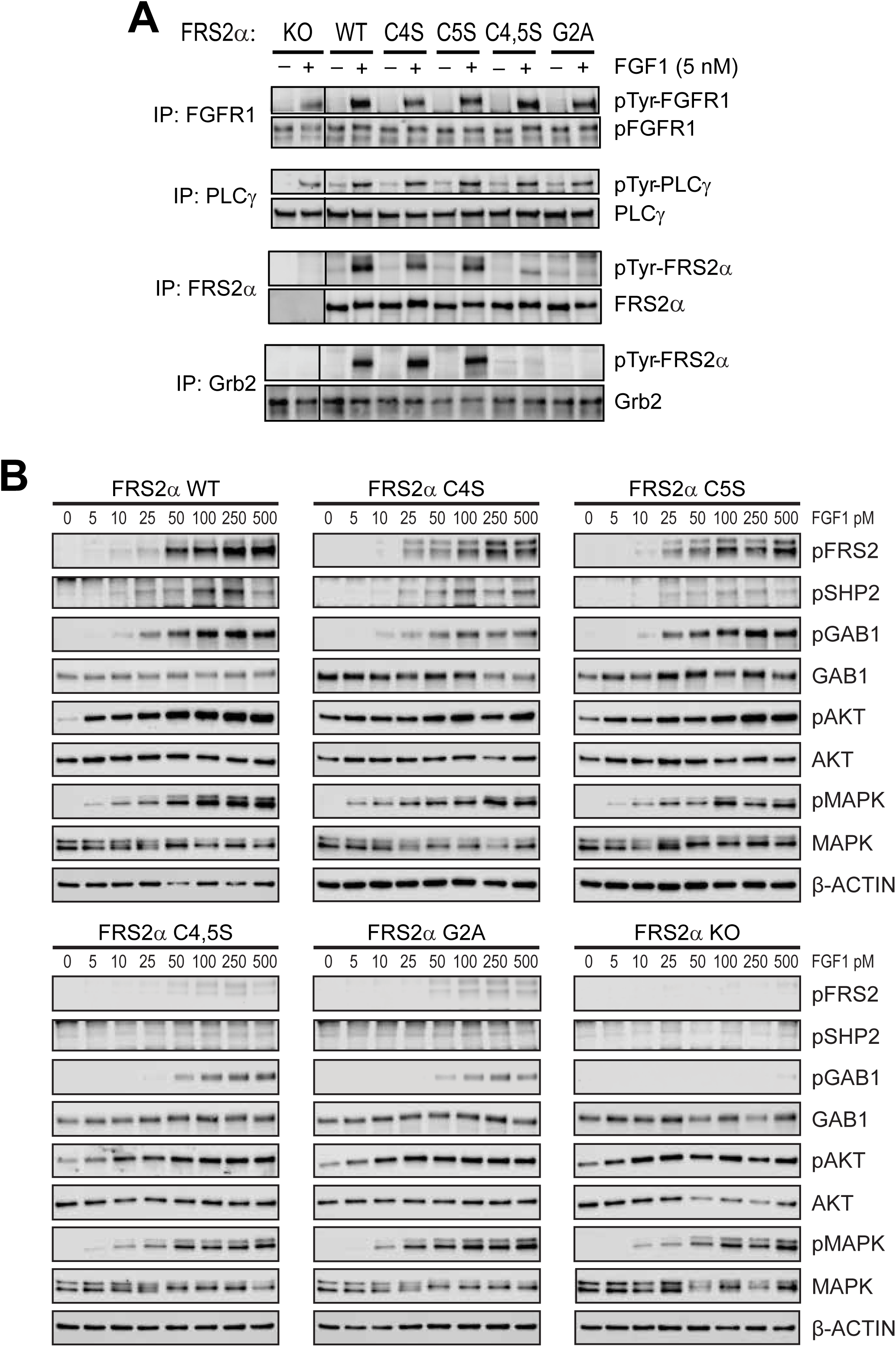
FGF1-induced signaling in FRS2α −/− MEFs and in cells rescued with FRS2α WT or its myristoylation and palmitoylation mutants. **(A)** FGF1 stimulation of tyrosine phosphorylation of FGFR1, PLCγ, and FRS2α in FRS2α−/− MEFs or in cells stably expressing WT, C4S, C5S, C4,5S, or G2A FRS2α mutants. Confluent FRS2α−/− MEFs and MEFs expressing FRS2α WT or the indicated FRS2α mutants were either left unstimulated or stimulated with 5 nM FGF1 for 10 min at 37°C. Cell lysates were immunoprecipitated overnight at 4 °C with anti-FGFR1, anti-PLCγ, anti-FRS2α, or anti-Grb2 antibodies, subjected to SDS-PAGE, and analyzed by immunoblotting with anti-phosphotyrosine (pTyr) antibodies to detect tyrosine phosphorylation. Immunoblotting with anti-FGFR1C, anti-PLCγ, anti-FRS2α, and anti-Grb2 antibodies was performed as a loading control. **(B)** Tyrosine phosphorylation of Gab1 and Shp2, as well as activation of AKT and MAPK, examined at increasing concentrations of FGF1 in FRS2α−/− MEFs and in cells expressing WT, C4S, C5S, C4,5S, or G2A FRS2α mutants. FRS2α−/− MEFs and MEFs expressing FRS2α WT or its myristoylation and palmitoylation mutants were left unstimulated or stimulated with increasing concentrations of FGF1 (5–500 pM) for 10 min at 37 °C. Cell lysates were subjected to SDS-PAGE and analyzed by immunoblotting for tyrosine phosphorylation of FRS2α, Shp2, and Gab1, as well as for activation of MAPK and AKT using anti-pFRS2α, anti-pShp2, anti-pGab1, anti-pMAPK, and anti-pAKT antibodies, respectively. Immunoblotting with anti-Gab1, anti-Shp2, anti-AKT, anti-MAPK, and anti-β-actin antibodies was performed as a loading control.

We have previously shown that the FRS2α G2A mutant is not tyrosine phosphorylated and, consequently, unable to recruit Grb2 in FGF1-stimulated cells^2^. Accordingly, we examined whether tyrosine phosphorylation of FRS2α depends on one or both palmitoylation sites. Lysates of FGF1-stimulated and untreated cells were immunoprecipitated with either anti-FRS2α or anti-Grb2 antibodies and analyzed for FRS2α tyrosine phosphorylation with anti-pTyr antibodies (Fig. 2A, third and fourth lanes). Anti-FRS2α and Anti-Grb2 antibodies were used as a loading control. We found that the single cysteine mutants FRS2α C4S and C5S, but not the palmitoylation deficient mutant FRS2α C4,5S and the myristoylation mutant G2A, could be tyrosine phosphorylated by FGF1 stimulation (Fig. 2A, third and fourth lanes). Immunoblotting with anti-FRS2α or anti Grb2 antibodies showed similar FRS2α and Grb2 expression levels for all cell lines (Fig. 2A, third and fourth lanes). Thus, these experiments demonstrate that partial membrane association conferred by a single palmitoylation site is sufficient for FGF1 induced tyrosine phosphorylation of FRS2α.

We wished to ascertain whether signaling pathways that are activated in response to FGF1 stimulation are affected by the palmitoylation state of FRS2α. For this purpose, we initially performed a detailed dose-response experiment of FGF1 (5–500 pM) to assess how changes in the palmitoylation state of FRS2α affect tyrosine phosphorylation of key substrates recruited via FRS2α to the membrane as well as activation of intracellular signaling pathways induced by FGF1 stimulation. These experiments also allowed us to determine the lowest concentration of FGF1 at which tyrosine phosphorylation of FRS2α WT reaches saturation. As shown in Figure 2B, tyrosine phosphorylation of Shp2 and Gab1 was virtually eliminated in FRS2α−/− MEFs across the full range of FGF1 concentrations. By contrast, AKT and MAPK remained robustly phosphorylated in FRS2α−/− MEFs stimulated with 50–500 pM of FGF1. These results indicate that tyrosine phosphorylation of Gab1 and Shp2 strictly depend on the integrity of FRS2α, whereas activation of MAPK and AKT in FRS2α-deficient cells can proceed via alternative compensatory signaling mechanisms.

These experiments also demonstrated that FRS2α−/− MEFs reconstituted with wild-type FRS2α show detectable Gab1 tyrosine phosphorylation at FGF1 concentrations as low as 25 pM (Fig. 2B). Also, FRS2α−/− MEFs expressing the C4S or C5S mutant exhibited strong and nearly complete restoration of Shp2 and Gab1 tyrosine phosphorylation across the full range of FGF1 concentrations (25–500 pM). Tyrosine phosphorylation of Gab1 was also restored in FRS2α−/− MEFs expressing the G2A or C4,5S mutant, when stimulated with FGF1 concentrations of 50–500 pM, albeit to a lower extent (Fig. 2B). Notably, weak tyrosine phosphorylation of Shp2 was also detectable in these cells at the higher range of FGF1 concentrations (100–500 pM), which may contribute to the increased Gab1 phosphorylation (Fig. 2B).

To further investigate how the palmitoylation state of FRS2α influences signaling dynamics downstream of FGF1, we next performed time-course experiments (Supplemental Fig 2). FRS2α−/− MEFs, along with MEFs expressing FRS2α WT or its lipidation mutants, were left unstimulated or stimulated with 0.1 nM FGF1, the concentration at which tyrosine phosphorylation of FRS2α WT and its effectors reaches saturation—for up to 6 h. As shown in the results, tyrosine phosphorylation of Shp2 was undetectable in FRS2α−/− MEFs and barely detected in cells expressing the C4,5S or G2A mutants. Tyrosine phosphorylation of Gab1 was also completely abolished in FRS2α−/− MEFs and was weak and transient in cells expressing the C4,5S or G2A mutants, disappearing after ∼15 min. By contrast, in MEFs expressing FRS2α WT or the single-site mutants C4S and C5S, Gab1 phosphorylation persisted for up to 1 h. Despite the pronounced defects in Shp2 and Gab1 phosphorylation in cells expressing the lipidation mutants, activation of both AKT and MAPK remained comparable across all cell lines. These findings, consistent with the dose-response experiments, indicate that MEFs rescued with lipidation-deficient FRS2α mutants acquire compensatory mechanisms that sustain AKT and MAPK activation, thereby supporting cell survival and proliferation in the absence of fully functional FRS2α.

Taken together, these experiments demonstrate that partial membrane association mediated by a single palmitoylation site on FRS2α is sufficient to support FGF1-induced tyrosine phosphorylation of the docking protein and to restore efficient MAPK and AKT signaling responses through assembly of Gab1 and other signaling molecules into complexes with tyrosine-phosphorylated FRS2α. Moreover, tyrosine phosphorylation of Gab1 can occur through both FRS2α-dependent and FRS2α-independent mechanisms. These findings also show that the MAPK and AKT pathways in FRS2α−/− MEFs have developed compensatory mechanisms for their activation that enable cell survival and proliferation despite the genetic loss of FRS2α.

### FGF1-induced membrane ruffling depends on the palmitoylation state of FRS2α

Initial visual inspection of confocal microscopy images of FRS2α WT-tdStayGold in FRS2α−/− MEFs or cells expressing FRS2α mutants revealed differences in membrane ruffling. It is well known that RTK ligands and other extracellular cues, induce membrane protrusions, for example, lamellipodia and growth cones during cell migration and neurite outgrowth, respectively, by altering the actin cytoskeleton^28^. To examine the role of FRS2α in actin remodeling, we used confocal microscopy to image FRS2α WT-tdStayGold in FRS2α−/− MEFs with or without FGF1 stimulation. In these experiments, we visualized FRS2α doubly via the fluorescence of tdStayGold and anti-Myc-tag staining (since FRS2α WT-tdStayGold possesses a C-terminal Myc-tag) and the actin cytoskeleton by phalloidin staining. We found that FGF1 (5 nM) induced the formation of peripheral membrane ruffles, which were identified by their irregular shape and intense phalloidin signal (Fig. 3A). Notably, these structures were enriched in FRS2α expressing cells. On the other hand, in unstimulated cells, there was little phalloidin staining resembling ruffles, and any FRS2α that could be seen at the cell periphery colocalized poorly with phalloidin. It should be noted that these experiments are shown as maximum intensity projections of confocal z-stacks, and as such, due to the flatness of the cells, the membrane localization of FRS2α is somewhat obscured; however, the presence of FRS2α in ruffles is conspicuous because these structures curl upwards in the z-axis.

**Figure 3.**
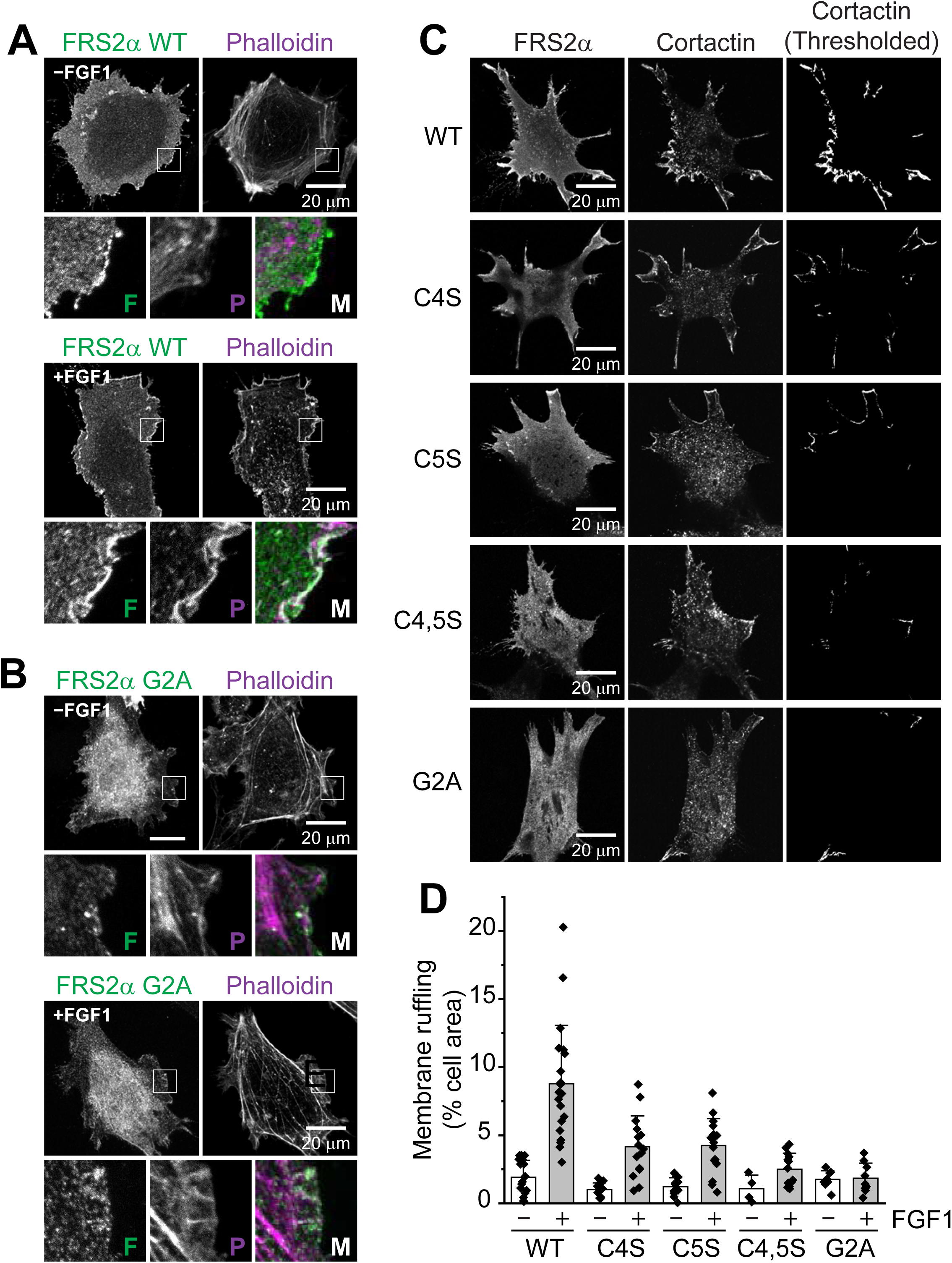
FGF1-induced membrane ruffling depends on the palmitoylation state of FRS2α. **(A, B)** Confocal images of FRS2α−/− MEFs transiently expressing FRS2α WT-tdStayGold (A) or FRS2α G2A-tdStayGold (B) with or without stimulation with FGF1 (5 nM) for 5 min at 37 °C. Cells were fixed and stained with phalloidin. Maximum intensity projections of z-stacks are shown. Magnified views of boxed regions are displayed in bottom three panels labeled “F” for FRS2α, “P” for phalloidin and “M” for merged. **(C)** Cortactin immunostaining of FGF1-stimulated FRS2α−/− MEFs expressing FRS2α WT-tdStayGold or its lipidation mutants fused to td-StayGold. Images are maximum intensity projections of bottom 3–4 confocal planes. Rightmost panels show thresholded cortactin immunofluorescence only at the plasma membrane. **(D)** Quantification of membrane ruffling, as a percentage of cell area, based on cortactin immunofluorescence at the plasma membrane. The extent of membrane ruffling for FRS2α WT is significantly different (P>0.001) from those of C4S and C5S, which in turn are significantly different (P>0.01) from those of C4,5S and G2A. Error bars indicate mean ± SE.

In contrast to the above results, FRS2α−/− MEFs expressing the G2A mutant did not show obvious membrane ruffling after FGF1 stimulation (Fig. 3B). Even membrane regions that were likely lamellipodia, for example, based on the presence of dorsal stress fibers that are perpendicular to the leading edge^29^ (magnified view “P” in lower panels of Fig. 3B), did not show the strong, irregularly shaped phalloidin staining expected of ruffles. Interestingly, although FRS2α G2A does not associate with the membrane, it appeared to colocalize very weakly with some stress fibers in FGF1-stimulated cells, possibly reflecting an interaction with factors associated with the actin cytoskeleton.

We wished to quantify membrane ruffling by FRS2α WT and its lipidation mutants. However, we deemed this to be a difficult task using phalloidin because of the complexity of the actin cytoskeleton. For example, in some cases, it would not be possible to unambiguously separate membrane ruffles from nearby peripheral stress fibers. For this reason, we turned to imaging cortactin, which is a protein involved in actin remodeling that associates with F-actin only at sites of dynamic actin assembly^30^ and thus can serve as a marker for membrane ruffles^31^. We transiently expressed FRS2α-tdStayGold constructs in FRS2α−/− MEFs, which were grown to low confluency (20–30%), and either stimulated the cells with FGF1 (5 nM) or left them untreated. We then performed immunostaining using an anti-cortactin antibody and imaged the cells by confocal microscopy. Cortactin fluorescence was thresholded with a fixed cutoff value and the fluorescence within the cell, which was punctate and likely represented the labeling of endosomal actin tails^30^, was cleared so that only the fluorescence of membrane ruffles remained for image analysis (Fig. 3C). In agreement with phalloidin staining experiments (Fig. 3A and 3B), cortactin immunostaining showed that FRS2α WT supported prominent FGF1-induced ruffling, while FRS2α G2A did not (Fig. 3C and 3D). This lack of stimulated ruffling was also observed in cells expressing FRS2α C4,5S. On the other hand, cells expressing the single palmitoylation mutants FRS2α C4S and C5S showed some stimulated ruffling that was significantly greater than that of the palmitoylation deficient mutants but less than that of FRS2α WT. Thus, these experiments demonstrate that robust FGF1-induced membrane ruffling requires both palmitoylation sites of FRS2α.

### FGF1 induces the formation of FRS2α-labeled macropinosomes

To further study the role of FRS2α in membrane ruffling, we performed live-cell, time-lapse confocal imaging of FRS2α WT-tdStayGold in FRS2α−/− MEFs. As shown in maximum intensity projections (Fig. 4A, upper panels), FGF1 (5 nM) induced membrane remodeling that was marked by a dramatic increase in FRS2α fluorescence at the leading edge of the cell, where membrane expansion occurred. Inspection of individual confocal planes revealed the formation of membrane ruffles strongly labeled with FRS2α, which was followed by the transient appearance of large vesicles with well-defined lumens (Fig. 4A, lower panels) that were also strongly labeled with FRS2α. Based on their spatiotemporal correlation with membrane ruffles and size (∼1 µm in diameter), these vesicles likely represented macropinosomes, as macropinocytosis is a form of fluid-phase endocytosis mediated by membrane ruffling^32^. Furthermore, it has been reported that FGF-dependent FGFR1 internalization in endothelial cells is dependent on macropinocytosis^33^.

**Figure 4.**
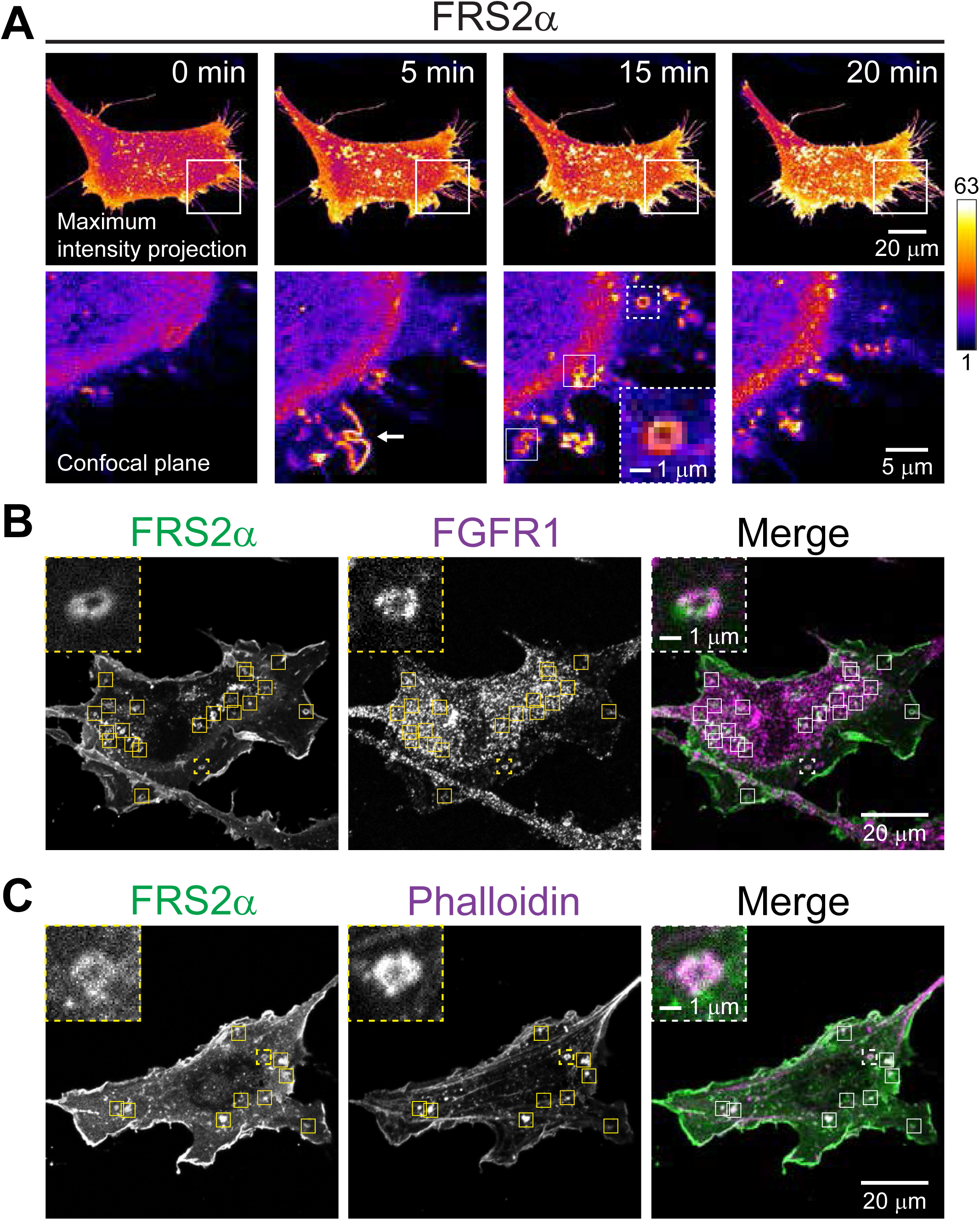
FGF1 promotes the formation of macropinosomes labeled with FRS2α, FGFR, and F-actin. **(A)** Timelapse confocal microscopy of FRS2α−/− MEFs transiently expressing FRS2α WT-tdStayGold stimulated with FGF1 (5 nM). Upper panels show a sequence of maximum intensity projections of z-stacks. Lower panels show sequence of magnified images of boxed regions in upper panels. Arrow and boxed regions in lower panels indicate the formation of membrane ruffles and macropinosomes, respectively. Inset shows magnified view of macropinosome in boxed region with dashed lines. Images are displayed using the “fire” lookup table in ImageJ. **(B, C)** Confocal images of FRS2α−/− MEFs expressing FRS2α WT-tdStayGold stimulated with FGF1 (5 nM), fixed and stained with anti-FGFR1 antibody **(B)** or phalloidin **(C)**. Maximum intensity projections of 2–3 middle confocal planes are shown. Insets show magnified views of macropinosomes in boxed regions with dashed lines.

Accordingly, we examined whether FRS2α-labeled vesicles harbor FGFR1, which should be the case if these vesicles are endocytic. FRS2α−/− MEFs transiently expressing FRS2α WT-tdStayGold were stimulated with FGF1, fixed and stained with FGFR1 antibody for confocal microscopy. We made maximum intensity projections of several planes near the base of the cell, which showed that FGFR1 colocalized with FRS2α-labeled vesicles (Fig. 4B). This was particularly evident near the cell periphery, at lamellipodial regions, where the density of FGFR1-positive spots was relatively low and did not obscure colocalization.

Macropinosomes are molded by actin polymerization and thus initially coated with F-actin; they lose their coat as they escape the lamella and enter the cell interior^34^. As such, we tested whether the FRS2α-labeled vesicles possess F-actin or not. FRS2α−/− MEFs transiently expressing FRS2α WT-tdStayGold were stimulated with FGF1, fixed and stained with phalloidin, and then confocal images were again examined as maximum intensity projections of several planes near the base of the cell. As expected, we found that phalloidin decorated FRS2α-labeled vesicles (Fig. 4C). Together, the above experiments strongly suggest that FRS2α-labeled vesicles represent macropinosomes.

### Macropinocytosis depends on both palmitoylation sites of FRS2α

Serum albumin undergoes macropinocytosis and subsequent trafficking to lysosomes, where it is degraded to provide nutrients for cells^35^. This macropinocytic process can be monitored by using a heavily BODIPY-dye-labeled albumin (DQ-BSA) whose fluorescence is initially self-quenched but, through lysosomal digestion, increases in brightness by severalfold^35,36^. To examine the role of FRS2α palmitoylation in FGF1-induced macropinocytosis, we performed the DQ-BSA dequenching assay with FRS2α−/− MEFs stably expressing FRS2α WT or its lipidation mutants. Cells grown to low confluency (20–30%) were stimulated with FGF1 (5 nM) or left untreated in media containing DQ-BSA. After extensive washing on ice, cells were chased for 2 h in fresh media to allow lysosomal degradation of DQ-BSA to occur, fixed and imaged by confocal microscopy. Images were then examined as maximum intensity projections. We found that DQ-BSA fluorescence was bright with FRS2α WT but very dim with the lipidation mutant cells, with the palmitoylation deficient mutants FRS2α C4,5S and G2A showing the least amount of fluorescence, comparable to that of unrescued FRS2α−/− MEFs (Fig. 5Ai and 5Aii). These experiments demonstrate that robust macropinocytosis requires both palmitoylation sites of FRS2α.

**Figure 5.**
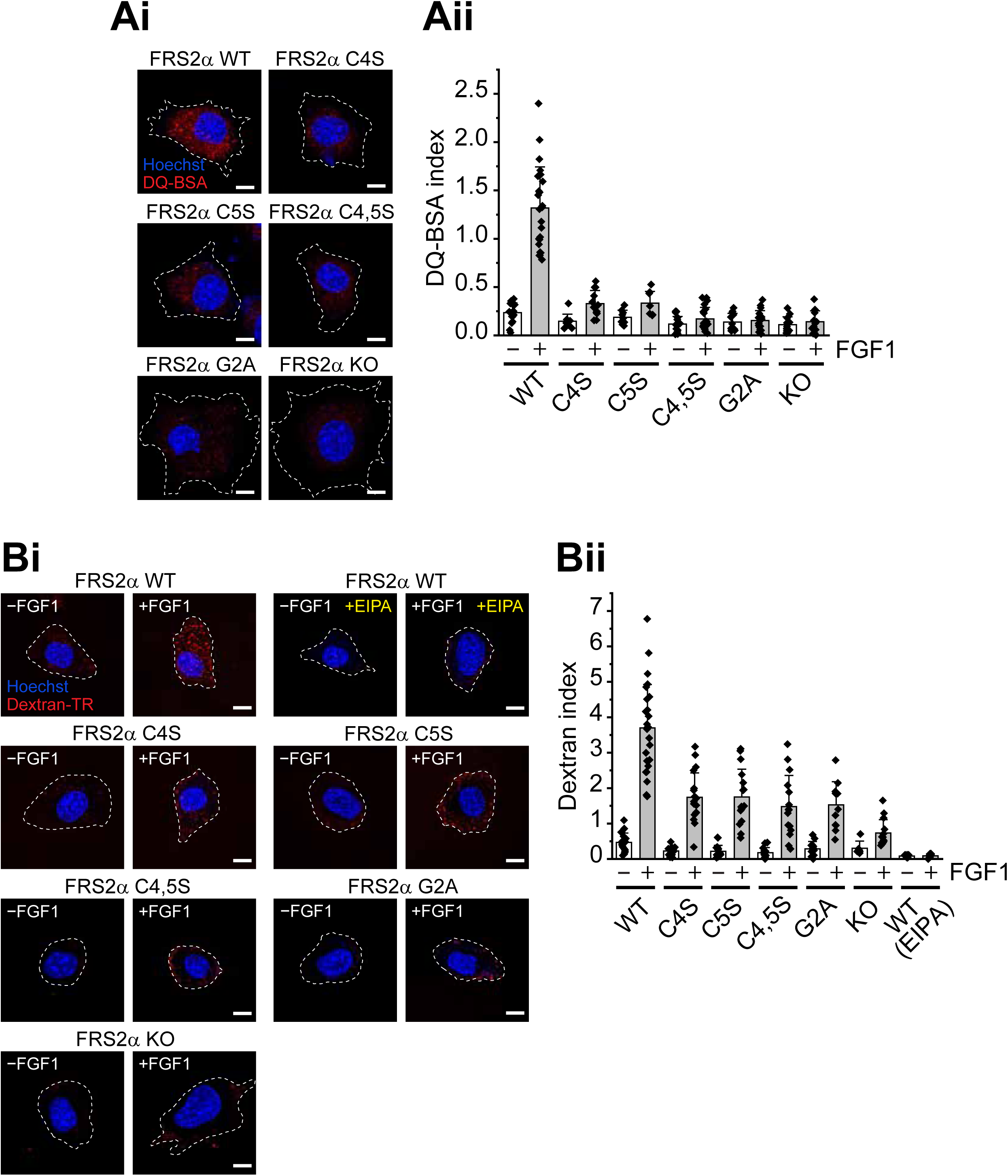
FGF1-induced macropinocytosis depends on FRS2α palmitoylation. **(Ai, Aii)** DQ-BSA dequenching assay with FRS2α−/− MEFs stably expressing FRS2α WT or its lipidation mutants. Confocal images of cells stimulated with FGF1 (5 nM) for 5 min at 37 °C or left untreated in media containing DQ-BSA and subsequently chased for 2h in fresh media **(Ai)**. Maximum intensity projections of z-stacks are shown. Quantification of DQ-BSA fluorescence are normalized to cell area **(Aii)**. DQ-BSA index of FGF1-stimulated cells are significantly different (P<0.001) than those of all mutants. Of note, DQ-BSA indices of FGF1-stimulated cells expressing C4,5S and G2A are not significantly different from that of FRS2α−/− (KO) cells. Error bars indicate mean ± SE. **(Bi, Bii)** Dextran uptake assay with FRS2α−/− MEFs stably expressing FRS2α WT or its lipidation mutants. Confocal images of cells stimulated with FGF (5 nM) for 5 min at 37 °C or left untreated in media containing Dextran-TR **(Bi).** To block macropinocytosis, cells were pretreated with 75 µM EIPA. Maximum intensity projections of z-stacks are shown. Quantification of dextran uptake normalized to cell area **(Bii)**. Dextran index of FGF1-stimulated cells is significantly different (P<0.001) than those of all mutants. Of note, dextran index of FGF1-stimulated cells expressing G2A is significantly different (P<0.001) from that of FRS2α −/− cells. Error bars indicate mean ± SE.

To corroborate these findings, we used another established macropinocytosis marker, high-molecular weight dextran. Unlike DQ-BSA, dye-labeled dextran is constitutively fluorescent, which allows macropinosomes to be tracked at early stages. We first confirmed that dextran and BSA undergo macropinocytosis together in FRS2α−/− MEFs stably expressing FRS2α WT by using Texas-Red labeled dextran (Dextran-TR) and fluorescein-labeled BSA (BSA-FITC), which is also constitutively fluorescent. Cells were stimulated with FGF1 in media containing both dextran-TR and BSA-FITC, fixed and imaged by confocal microscopy. As expected, the two cargos showed extensive colocalization, indicating that they were similarly internalized by macropinocytosis (Supplemental Fig. 3).

Next, FRS2α−/− MEFs that stably express FRS2α WT or its lipidation mutants were grown to low confluency (20–30%) and stimulated with FGF1 (5 nM) or left untreated in media containing dextran-TR. After extensive washing on ice, cells were fixed and imaged by confocal microscopy. Images were then examined as maximum intensity projections. We found that FGF1-induced dextran-TR uptake was robust in cells expressing FRS2α WT and very weak in FRS2α−/− MEFs, but that it was not as diminished as we had expected in cells expressing the lipidation mutants (Fig. 5Bi). Notably, while dextran-TR spots were spatially distributed throughout the cell with FRS2α WT, they were restricted to the cell periphery with the lipidation mutants, particularly with the palmitoylation deficient mutants FRS2α C4,5S and G2A. Importantly, dextran-TR staining was eliminated in cells pretreated with EIPA, an inhibitor of micropinocytosis^37^.

To quantify dextran-TR uptake, we measured the percentage of cell area occupied by dextran-TR-positive macropinosomes as previously described^38,39^, to calculate the “macropinocytic index.” This analysis indicated that while stimulated uptake of dextran-TR was significantly reduced with the lipidation mutants compared to FRS2α WT, it was indeed higher than expected (Fig. 5Bii). Specifically, in the DQ-BSA dequenching assay, the macropinocytosis signal for FRS2α C4,5S and G2A in stimulated cells was only ∼13% and ∼12% compared to FRS2α WT, respectively. However, in the dextran-TR uptake assay, the analogous macropinocytosis signal for FRS2α C4,5S and G2A was ∼39% and ∼40%, respectively. Given that this relatively mild reduction in stimulated dextran-TR uptake is correlated with the spatial restriction of dextran-TR spots at the cell periphery, we wondered if FGF1-induced macropinocytosis is defective at an early step with the palmitoylation deficient FRS2α mutants. In comparison, with the single cysteine mutants, the FGF1-induced macropinocytosis signal obtained with DQ-BSA and dextran-TR did not differ as much: They were ∼31% and ∼40%, respectively, which suggested that macropinocytosis was not as defective when FRS2α possessed at least one palmitoylation site.

### Macropinosome closure requires FRS2α palmitoylation

The mechanisms underlying macropinocytosis involve multiple steps that ultimately lead to the closure of membrane ruffles into macropinosomes^40,41,42,43^. Moreover, even after successfully sealing, nascent macropinosomes must leave the actin-dense lamellar region, by ostensibly deforming and squeezing through the actin cytoskeleton^34,44^, to fuse with lysosomes in the cell interior so that internalized solutes are degraded. Thus, defective macropinosome closure or transport through the lamella could explain the observed confinement of dextran-TR spots to the cell periphery with the palmitoylation deficient FRS2α mutants C4,5S and G2A.

Accordingly, we sought to determine which step of FGF1-induced macropinocytosis is affected when FRS2α palmitoylation is blocked. We chose to focus on macropinosome closure since there is precedent for this step being defective under certain conditions. For example, it has been observed that macropinosomes can form but do not seal when phosphatidylinositol 3-kinase is inhibited^40,42^.

One way to potentially assay macropinosome closure is to test the accessibility of dextran-TR-labeled macropinosomes—or perhaps more accurately, membrane ruffling structures—in intact cells to extracellularly applied anti-dextran antibodies. To this end, FRS2α−/− MEFs that stably express FRS2α WT or the palmitoylation deficient mutant FRS2α G2A were subjected to the dextran uptake assay, this time with a subsequent anti-dextran immunostaining step. Because parental FRS2α−/− MEFs, without any rescue, showed the least amount of dextran uptake, they were also studied. As in previous experiments, cells grown to low confluency (20–30%) were stimulated with FGF1 in media containing dextran-TR. After extensive washing on ice, cells were fixed (without permeabilization) and sequentially stained with anti-dextran antibodies and Alexa488-conjugated secondary antibodies. Cells were then imaged by confocal microscopy and maximum intensity projections of dextran-TR and anti-dextran immunostaining were examined. We found that FRS2α WT cells, while showing strong dextran-TR uptake, were weakly stained with anti-dextran antibody, with very few dextran-TR spots colocalizing with the antibody-labeled spots (Supplemental Fig. 4). On the other hand, FRS2α G2A cells and parental FRS2α−/− MEFs both showed strong anti-dextran immunostaining with robust colocalization of dextran-TR and anti-dextran antibody spots at the cell periphery. Evidently, macropinocytic structures can retain dextran-TR without sealing in cells lacking FRS2α or when FRS2α is palmitoylation deficient. Thus, these experiments demonstrate that FGF1-induced macropinocytosis depends on FRS2α palmitoylation not only for membrane ruffling, but also for macropinosome closure.

### Neuronal differentiation of FGF1 stimulated PC12 cells depends upon FRS2α palmitoylation

The experiments described thus far were performed using fibroblasts derived from murine FRS2α−/− embryos (MEFs). Our next goal was to investigate the role of FRS2α palmitoylation in a different cellular context, specifically to explore its contribution to FGF1-induced cell differentiation. To this end, we examined FRS2α palmitoylation in neuronal differentiation of PC12 cells, a well-established model derived from rat pheochromocytoma. PC12 cells provide a robust system for studying the molecular mechanisms of neuronal differentiation induced by FGF stimulation of FGFRs, as well as by neurotrophic factors that signal through TrkA, TrkB, and RET receptor tyrosine kinases.

To explore the role of FRS2α palmitoylation in regulating cellular signaling pathways in PC12 cells, we generated FRS2α deficient PC12 cells using the CRISPR-Cas9 system and confirmed the absence of FRS2α expression in FRS2α−/− PC12 cells (Fig 6A). Similar to FRS2α−/− MEFs, FRS2α−/− PC12 cells are able to proliferate despite the loss of FRS2α expression. Parental PC12 cells, FRS2α−/− PC12 cells and FRS2α−/− PC12 cells rescued with FRS2α WT were left unstimulated or stimulated with FGF1 for 10 min at 37°C. Cell lysates were then immunoprecipitated with anti-FRS2α antibodies, subjected to SDS-PAGE, and analyzed by immunoblotting with anti-pTyr antibodies to determine the level of tyrosine phosphorylation of FRS2α or with anti-FRS2α antibodies to quantitate the expression level of FRS2α in these cells. This experiment confirmed that FRS2α−/− PC12 cells are indeed deficient in FRS2α and demonstrated a comparable expression level of ectopically expressed FRS2α WT in FRS2α−/− PC12 cells relative to the level of endogenous FRS2α in parental PC12 cells.

**Figure 6.**
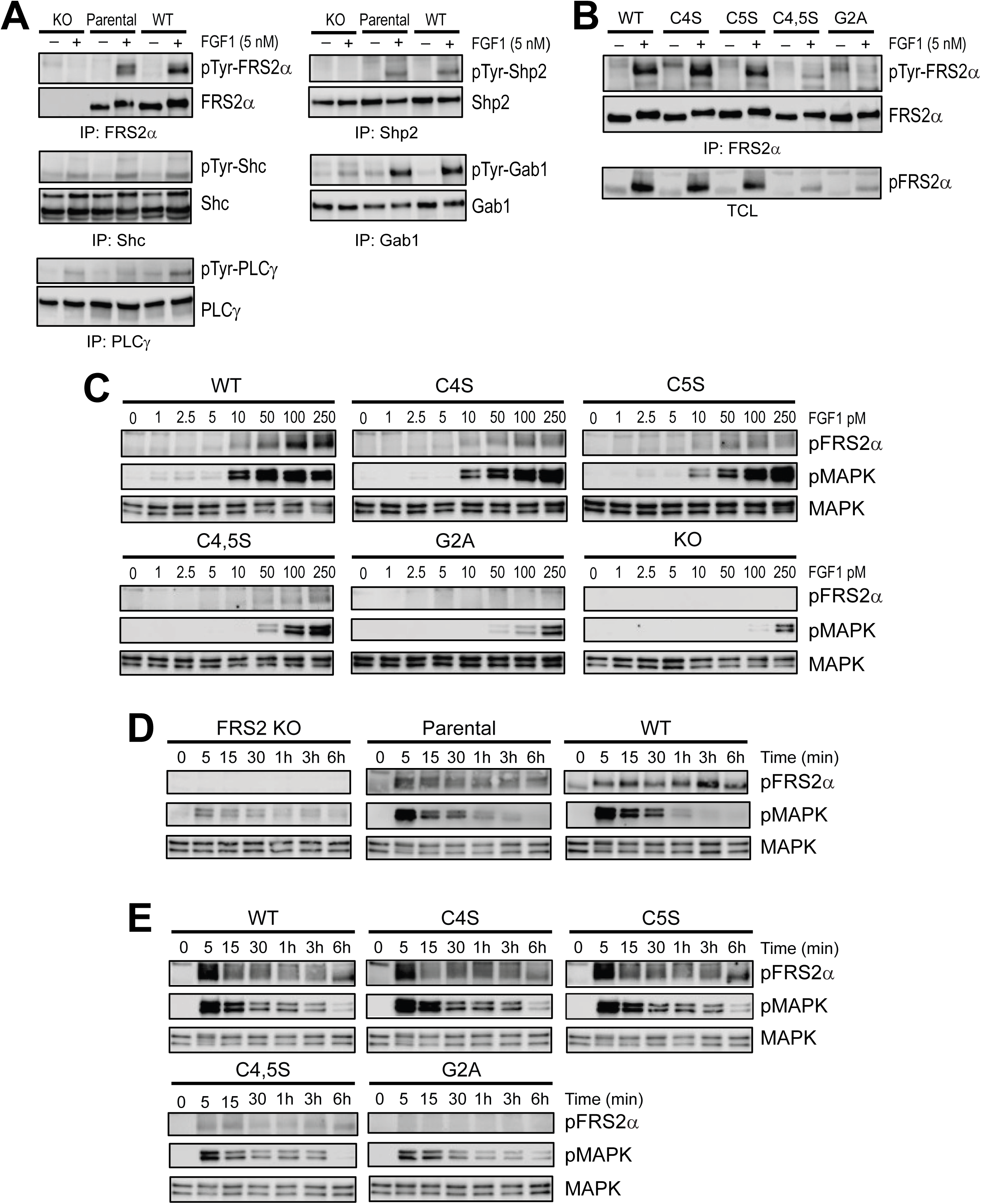
FGF1-induced signaling in FRS2α−/− PC12 cells and in cells rescued with FRS2α WT or its lipidation mutants. **(A)** Tyrosine phosphorylation of FRS2α, PLCγ, Shc, Shp2, and Gab1 in FRS2α-deficient and rescued PC12 cells expressing FRS2α WT, C4S, C5S, C4,5S, or G2A mutants. Confluent parental PC12 cells, FRS2α−/− (KO) PC12 cells, or FRS2α−/− PC12 cells rescued with FRS2α WT were either left unstimulated or stimulated with 5 nM FGF1 for 10 min at 37 °C. Cell lysates were immunoprecipitated overnight at 4 °C with antibodies against FRS2α, PLCγ, Shc, Shp2, or Gab1, subjected to SDS-PAGE, and analyzed by immunoblotting with anti-phosphotyrosine (pTyr) antibodies. Antibodies specific for each protein were used as loading controls. **(B)** FGF1 stimulation of tyrosine phosphorylation of FRS2α in FRS2α−/− PC12 cells expressing FRS2α WT or its lipidation mutants. Cells were left unstimulated or stimulated with 5 nM FGF1 for 10 min at 37 °C. Cell lysates were then immunoprecipitated overnight at 4 °C with anti-FRS2α antibodies and subjected to SDS-PAGE, followed by immunoblotting with anti-pTyr antibodies. Anti-FRS2α antibodies were used as a loading control (Fig.6B upper lane). Total cell lysates from the indicated cell lines were also analyzed by immunoblotting with anti-phospho-FRS2α antibodies (pFRS2α, Fig.6B lower lane). **(C)** Tyrosine phosphorylation of FRS2α and MAPK activation analyzed at increasing concentrations of FGF1 in FRS2α−/− PC12 cells left unrescued or rescued with FRS2α WT or its lipidation mutants. Cells were left unstimulated or stimulated with increasing concentrations of FGF1 (1–250 pM). Total cell lysates were subjected to SDS-PAGE and immunoblotted with anti-pFRS2α and anti-pMAPK antibodies. Immunoblotting with anti-MAPK antibodies was used as a loading control. **(D, E)** Kinetics of FRS2α and MAPK phosphorylation in FRS2α−/− PC12 cells left unrescued or rescued with FRS2α WT or its lipidation mutants. Cells were stimulated with FGF1 (0.25 nM) for the indicated times. Total cell lysates were subjected to SDS-PAGE and immunoblotted with anti-pFRS2α and anti-pMAPK antibodies. Anti-MAPK antibodies were used as loading controls.

We next investigated FGF1-induced tyrosine phosphorylation of downstream effectors proteins by immunoprecipitating cell lysates with antibodies against Shc, PLCγ, Shp2 or Gab1 followed by immunoblotting analysis (Fig. 6A). To assess the level of tyrosine phosphorylation of these proteins we used anti-pTyr antibodies and, as shown in Figure 6A, tyrosine phosphorylation of FRS2α and Shp2 was virtually eliminated in the knockout cells while tyrosine phosphorylation of Gab1 was markedly reduced. In contrast, tyrosine phosphorylation of Shc and PLCγ remained intact, indicating that their recruitment to activated FGFR1 and tyrosine phosphorylation induced by FGF1 stimulation is independent of FRS2α.

To determine how alterations in the palmitoylation state of FRS2α affect its tyrosine phosphorylation, lysates from untreated and FGF1-stimulated PC12 cells expressing either FRS2α WT or its various mutants were immunoprecipitated with anti-FRS2α antibodies, followed by immunoblotting with anti-pTyr antibodies. As shown in Figure 6B (upper lane), FRS2α WT and the single palmitoylation mutants C4S and C5S exhibited similar levels of tyrosine phosphorylation, whereas the double mutant C4,5S and the G2A mutant were barely phosphorylated on tyrosine. Anti-FRS2α immunoblotting was used as a loading control. Similar results were obtained using phospho-specific FRS2α antibodies (pFRS2α, lower panel).

We next investigated how alterations in the palmitoylation state of FRS2α affect the activation of MAPK response in PC12 cells by analyzing dose-dependent FGF1 responses in the range of 1–250pM in these cells (Fig 6C). These experiments also established the lowest FGF1 concentration at which tyrosine phosphorylation of FRS2α reaches saturation in PC12 cells. As expected, tyrosine phosphorylation of FRS2α was undetectable in FGF1-stimulated FRS2α−/− PC12 cells, and MAPK activation in these cells was only observed at 250 pM of FGF1. In G2A expressing PC12 cells, tyrosine phosphorylation of the G2A mutant was barely detectable with a weak MAPK response detectable at 50 pM which gradually increased at 100 and 250 pM of FGF1. In FGF1-stimulated C4,5S PC12 cells, tyrosine phosphorylation of the C4,5S mutant was weak at 50 pM, and slightly increased at 100 and 250 pM of FGF1 stimulation. The MAPK response in C4,5S PC12 cells was more pronounced, showing progressively stronger activation across the 50–250 pM FGF1 range. The profile of FGF1-stimulated tyrosine phosphorylation of WT, C4S, and C5S expressed in FRS2α−/− PC12 cells showed a similar gradual increase across the 10–250 pM FGF1 range, with FRS2α WT exhibiting more robust tyrosine phosphorylation compared to the singly palmitoylated C4S or C5S mutants. Similarly, the dose response of MAPK activation demonstrated a progressive increase from 10 to 250 pM FGF1, with enhanced MAPK phosphorylation in FRS2α WT-expressing PC12 cells relative to C4S- or C5S-expressing cells.

Measurements of the kinetics of FRS2α tyrosine phosphorylation and the activation of MAPK in response to FGF1 stimulation revealed similar kinetics for both parental PC12 cells and FRS2α−/− PC12 cells ectopically expressing FRS2α WT (Fig. 6D). In these cells, tyrosine phosphorylation of FRS2α WT and MAPK activation occurred rapidly within 5 min followed by a gradual decline between 15 min to 6 h. FGF1 stimulation of MAPK activation in FRS2α−/− PC12 cells also revealed rapid onset after 5 min followed by gradual decease in activation from 15 min to 6 h. In the absence of FRS2α expression, the MAPK response of PC12 cells was weak and entirely mediated via alternative cell signaling pathways that are activated by FGF1 stimulation of FGFR at the cell membrane (Fig. 6D).

The kinetics of tyrosine phosphorylation of FRS2α-deficient PC12 cells expressing the G2A mutant revealed barely detectable tyrosine phosphorylation of this mutant while the kinetics of PC12 cells expressing the C4,5S mutant revealed early stimulation at 5 to 15 min followed by gradual decline from 30 min to 6 h (Fig. 6E). MAPK activation in both the C4,5S and the G2A expressing cells revealed a peak of activity after 5 min followed by gradual decline over 6 h. The overall profiles of the kinetics of tyrosine phosphorylation and MAPK response in FRS2α−/− PC12 cells expressing either FRS2α WT or the C4S or C5S mutants revealed similar rapid onset of activation after 5 min followed by gradual declines. The kinetics experiments showed a weak MAPK response in cells expressing the G2A and C4,5S mutants with a steep decline relative to cells expressing FRS2α WT and the two singly palmitoylated FRS2α mutants. These results demonstrate that the robustness of MAPK signaling is compromised in the absence of palmitoylation of FRS2α (Fig. 6E).

We next explored the impact of FRS2α deficiency on FGF1-induced neuronal differentiation in parental PC12 cells, FRS2α−/− PC12 cells or FRS2α−/− PC12 cells ectopically expressing FRS2α WT. Cells were either left unstimulated or stimulated with 5 nM FGF1 for 48 h at 37°C, after which microscopic images of neuronal differentiation were obtained and neurite length quantified. The experiment shown in Figures 7A and B reveals robust FGF1-induced neuronal differentiation in both parental PC12 cells and FRS2α−/− PC12 reconstituted with FRS2α WT, but not in FRS2α−/− PC12 cells. To explore the role played by palmitoylation in FGF1 stimulation of neuronal differentiation, similar experiments were performed with PC12 cells ectopically expressing FRS2α WT, C4S, C5S, C4,5S or the G2A mutant. The experiment shown in Figures 7C and D reveals robust FGF1 stimulation of neuronal differentiation with FRS2α WT, C4S and C5S ectopically expressed in FRS2α−/− PC12 cells, weak neuronal differentiation with C4,5S expressed in FRS2α−/− PC12 cells, and no neuronal differentiation with G2A expressed in FRS2α−/− PC12 cells.

**Figure 7.**
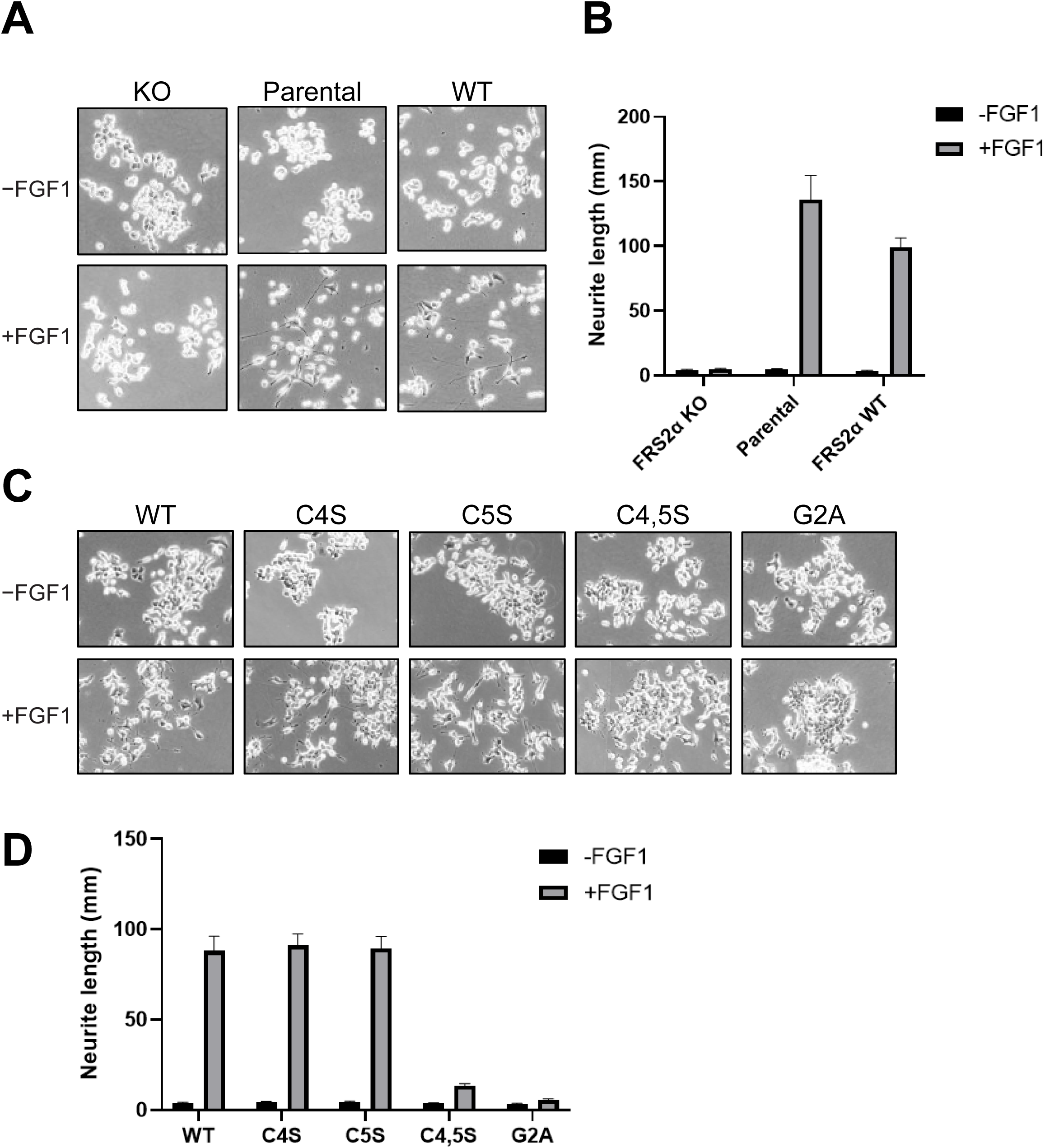
FGF1-induced neurite outgrowth in FRS2α−/− PC12 cells and in cells rescued with FRS2α WT or its lipidation mutants. (**A, C)** Neurite outgrowth assay. FRS2α−/− PC12 cells, parental PC12 cells, and FRS2α−/− PC12 cells rescued with FRS2α WT or its lipidation mutants were seeded in 10-cm plates (5 × 10^5^ cells per plate). After 24 h, cells were stimulated with 5 nM FGF1, and neurite outgrowth was visualized after 72 h of incubation. Representative microscopic images are shown. **(B, D)** Quantification of neurite length (unstimulated black bars, FGF1stimulated gray bars). The average neurite length was measured and quantified for all cell lines as indicated. Data represent mean ± SD from three independent experiments.

Taken together, these experiments clearly establish that the integrity of FRS2α is essential for FGF1-induced neuronal differentiation of PC12 cells and that occupation of a single palmitoylation site of FRS2α is sufficient for efficient stimulation of intracellular signaling pathways required for FGF1-induced neuronal differentiation.

## Discussion

The docking protein FRS2α was discovered more than two decades ago as a central molecular hub of cellular signaling pathways activated at the cell membrane by ligand binding to FGFRs, NGFRs, and other RTKs stimulated by neurotrophic factors^2,3^. It was also shown that membrane association of FRS2α via myristoylation is essential for complex formation with either unstimulated RTKs (e.g., FGFRs) or ligand-activated RTKs (e.g., NGFRs). This association enables tyrosine phosphorylation of FRS2α in response to ligand binding to the extracellular domain, RTK dimerization, and activation of their intrinsic catalytic activity. It is well established that myristoylation of a variety of enzymes and proteins is essential for membrane binding and for eliciting their activities at the cell membrane. However, quantitative binding experiments have demonstrated that the dissociation constant (Kd) of a myristoyl moiety for the membrane^45^ is approximately 0.1 mM. Stable membrane association is therefore typically accomplished by an additional membrane targeting signal. Membrane binding can be reinforced by increasing the protein’s hydrophobicity through palmitoylation of cysteine residues, which complements myristoylation. Alternatively, a short stretch of positively charged amino acids that interact with negatively charged phospholipid headgroups on the inner leaflet of the membrane enhances the stability of membrane association^45^.

A more recent study showed that, besides myristoylation, FRS2α also undergoes palmitoylation^18^. Palmitoylation is mediated by a reversible reaction in which palmitate is attached to and released from cysteine residues via a thioester linkage, catalyzed by specific acylation and deacylation enzymes^19^. Stable association of FRS2α with the cell membrane is mediated by myristoylation and palmitoylation, and this is further supported by binding of the PTB domain of FRS2α to a specific binding site in the juxtamembane domain of FGFR1 and other members of the family. Because the palmitoylation of two cysteine residues in the N-terminal region of FRS2α is a reversible process, and membrane association of FRS2α is critical for mediating intracellular signaling, it is plausible that cellular responses to FGFs, NGF, and other neurotrophic factors can be regulated by FRS2α palmitoylation.

The cellular localization of FRS2α was explored in this study by using expression vectors encoding either wild-type FRS2α or mutants deficient in myristoylation or palmitoylation, each fused to tdStayGold. The fluorescently tagged FRS2α variants were expressed in FRS2α−/− MEFs, and their subcellular localization was visualized and quantified by confocal fluorescence microscopy. These experiments confirmed that fluorescently labeled FRS2α WT is predominantly localized at the plasma membrane, whereas both the palmitoylation-deficient (C4,5S) and myristoylation-deficient (G2A) mutants failed to associate with the cell membrane. In contrast, the singly palmitoylated mutants (C4S or C5S) exhibited reduced, but detectable, membrane association, corresponding to approximately 39% and 25% of the membrane fluorescence intensity observed with FRS2α WT, respectively.

We have previously demonstrated that MAPK activation and cell proliferation induced by physiological concentrations of FGF1 (10–50 pM) were markedly reduced in FRS2α−/− MEFs^12^. However, both MAPK signaling and cell proliferation were observed when these FRS2α-deficient MEFs were stimulated with high, non-physiological concentrations of FGF1 (10 nM). These findings indicate that at high ligand concentrations, FGF1 can bypass the requirement for FRS2α by activating alternative signaling pathways downstream of FGFRs. It is noteworthy that EGF- and serum-induced proliferation of MEFs are unaffected by the absence of FRS2α.

The experiments described in this manuscript demonstrate that palmitoylation of FRS2α plays a critical role in FGFR-induced cell signaling by regulating both the robustness and magnitude of tyrosine phosphorylation of FRS2α. Additional key conclusions of this manuscript are:

- Membrane association of FRS2α is regulated by palmitoylation of the docking protein.
- FGF-induced MAPK activation and other cellular responses are partially restored in cells expressing FRS2α mutants deficient in either one of the two FRS2α palmitoylation sites.
- In addition to controlling the strength of signaling, the anchoring of FRS2α to the inner leaflet of the plasma membrane provides a platform and organizational hub for the assembly of multi-protein complexes that are essential for cytoskeletal change associated with membrane ruffling, macropinocytosis and other FGF1-induced cellular processes. We also demonstrate that macropinocytosis requires palmitoylation of both sites of FRS2α.
- While FRS2α−/− PC12 cells are able to proliferate, FGF1 induced neuronal differentiation of these cells depends upon palmitoylation of FRS2α.

Intracellular signaling pathways activated by FGFRs and other RTKs are subject to multiple stimulatory cues as well as diverse negative feedback mechanisms operating within the same pathways. For example, FRS2α-mediated activation of the MAPK pathway, which is essential for many cellular responses, also initiates negative feedback regulation^10^. It has been shown that MAPK, when stimulated by FGF1 or by other extracellular ligands such as EGF, PDGF, LPA, or carbachol, phosphorylates FRS2α on eight threonine residues. This phosphorylation attenuates the MAPK response by interfering with the formation of signaling complexes between Grb2 and tyrosine-phosphorylated FRS2α.

The experiments presented in this report demonstrate that cellular signaling pathways mediated by FRS2α are regulated by the extent of palmitoylation of this docking protein. Protein palmitoylation is enzymatically controlled by protein acyl transferases, which catalyze the acylation of FRS2α, and by acyl-protein thioesterases and palmitoyl-protein thioesterases, which catalyze its de-acylation^19^. Accordingly, the level of FRS2α palmitoylation can be modulated by regulating the activities of these acylating and/or de-acylating enzymes. It is tempting to speculate that MAPK or other downstream components of the signaling pathways activated by FRS2α may also participate in a feedback loop that fine-tunes the palmitoylation state of FRS2α (Supplemental Fig. 5).

## Materials and Methods

### Plasmids, antibodies, growth factors, and reagents

The viral expression vector for FGFR1C was previously described^46^ and was generated by subcloning FGFR1C into pBABE-puro. Viral expression vectors for FRS2α WT and its lipidation mutants (C4S, C5S, C4,5S, and G2A) were generated by subcloning the corresponding FRS2α cDNAs with a C-terminal Myc tag into either pBABE-hygro (for infection of FRS2α−/− MEFs) or pLenti-hygro (for infection of FRS2α−/− PC12 cells). For transient expression in FRS2α−/− MEF, cDNAs encoding human FRS2α WT and the various mutants (G2A, C4S, C5S, and C4,5S) with a C-terminal tdStayGold-Myc tag were subcloned into a modified pOptiVec vector (Invitrogen) from which the IRES and DHFR sequences were deleted.

Antibodies against FGFR1C were raised against the 18 C-terminal amino acids of the protein. Anti-FRS2α antibodies used for immunoprecipitation were a mixture of antibodies raised against the PTB domain and the C-terminal region of FRS2α expressed as GST-fusion proteins. Anti-Shc antibodies were raised against the PTB domain of Shc expressed as a GST-fusion protein. Commercial antibodies were obtained as follows: anti-phospho-Tyr-FRS2α (AF5126) anti-FRS2α (MAB4069), and anti-phospho-Shp2 (AF3790) (R&D Systems); anti-Gab1 (3232S), anti-phospho-Gab1 (3231S), anti-Akt (9272S), anti-phospho-Akt (4060S), anti-MAPK (4695S), anti-phospho-MAPK (4370S), anti-Rab5 (C8B1), anti-Rab7 (D95F2), anti-Rab11 (D4F5), anti-FGFR1 (D8E4), and Myc-tag (9B11) (Cell Signaling Technology); anti-β-actin (SC-47778, Santa Cruz Biotechnology); anti-cortactin (PA5-27134; Invitrogen); anti-dextran (60026; StemCell Technologies). Imaging reagents were obtained as follows: CellMask Deep Red (C10046), Phalloidin Alexa568 (A12380), Hoechst 33342 (H3570), DQ Red BSA (D12051), and Dextran-Texas Red, 70,000 MW, Lysine Fixable (D1864) (Invitrogen). Expression and purification of FGF1 was performed as previously described^47^.

### Cell culture

HEK 293 cells were maintained in Dulbecco’s Modified Eagle’s Medium (DMEM; Gibco) supplemented with 10% fetal calf serum (FCS; Gibco) and 1% penicillin/streptomycin (PS; Gibco). FRS2α knockout (FRS2α−/−) MEFs were previously described^12^. Cells were maintained in DMEM supplemented with 10% FCS and 1% PS. MEFs rescued with FRS2α WT or its lipidation mutants were cultured in the same medium supplemented with 100 μg/ml hygromycin B (Invitrogen). PC12 cells were obtained from ATCC and cultured in DMEM supplemented with 10% fetal bovine serum (FBS), 10% horse serum (HS), and 1% PS.

To generate FRS2α knockout PC12 cells (FRS2α−/− PC12), cells were infected with lentivirus produced from the eSpCas9-LentiCRISPR v2 vector (GenScript) containing Cas9 and a guide RNA targeting rat FRS2α (5′-AGTCATAGCCATATCGTCGT-3′), which disrupts Arg63 within the PTB domain. Lentivirus for PC12 infection were produced in HEK293T cells using third-generation packaging plasmids. Infected PC12 cells were selected with 1 μg/ml puromycin (Gibco) and then subjected to single cell cloning. Gene disruption was validated by sequencing the target locus and by immunoblotting for FRS2α protein.

FRS2α−/− PC12 cells were maintained in the same medium as parental PC12 cells with 1 μg/ml puromycin. FRS2α−/− PC12 cells rescued with FRS2α WT or lipidation mutants were cultured under identical conditions, with the addition of 100 μg/ml hygromycin B.

### CellMask staining

FRS2α−/− MEFs were plated onto 35-mm glass-bottom dishes (P35G-1.5-14-C, MatTek) at a seeding density of 45,000 cells per dish in phenol red-free DMEM (21063029, Gibco) containing 10% FBS and reverse transfected with 1 μg FRS2α-tdStayGold constructs using Lipofectamine 3000 reagent (L3000008, Invitrogen), according to the manufacturer’s instructions. Twenty-hours later, cells were washed twice in phenol red-free DMEM, incubated with CellMask Deep Red (0.5 μL stain in 1 mL media) for 5 min at room temperature, washed three times in DMEM and subjected to hypotonic solution composed of 4:1 (v/v) diH2O/media. Cells were quickly imaged before appreciable endocytosis of CellMask occurred. Confocal microscopy was performed with a Zeiss LSM 880 Airyscan confocal microscope using a 63x/1.4 oil objective.

### Immunofluorescence

Twenty-hours after reverse transfection with tdStayGold constructs, FRS2α−/− MEFs were washed twice with phenol red-free DMEM and then incubated with or without FGF1 (5 nM) for 5 min at 37°C. Cells were then fixed in 4% paraformaldehyde (PFA; 15710, Electron Microscopy Sciences) for 15 min at room temperature, washed in PBS three times for 5 min, incubated for 1 h in blocking buffer (1x PBS, 5% normal goat serum, 0.3% Triton X-100), and then incubated overnight at 4 °C with primary antibody in antibody dilution buffer (1x PBS, 1% BSA / 0.3% Triton X-100) using the following antibody dilutions: 1:200 for anti-Rab5), 1:50 for anti-Rab7, 1:50 for anti-Rab11, 1:1500 for anti-Myc-tag, 1:500 for anti-cortactin, 1:500 for anti-FGFR1 and 1:100 for anti-dextran antibody. Cells were then washed in PBS three times for 5 min, incubated for 1 h with Alexa488- or Alexa568-conjugated goat anti-mouse or anti-rabbit highly cross-absorbed secondary antibody (Invitrogen) in blocking buffer, and washed in PBS three times for 5 min. Cells were imaged by confocal microscopy.

For membrane ruffling experiments based on cortactin immunostaining, cells were investigated at low (20–30%) confluence to avoid intercellular adhesion. Confocal images of isolated cells were examined as maximum intensity projections of the bottom 2–3 confocal planes which were thresholded with uniform threshold values across conditions.

### Macropinocytosis imaging and quantification

Macropinocytosis was assayed using DQ-BSA and dextran-TR as described^35^. For DQ-BSA experiments, FRS2α−/− MEFs stably expressing FRS2α WT or its lipidation mutants were plated onto 35-mm glass-bottom dishes (P35G-1.5-14-C, MatTek) at a seeding density of 45,000 cells per dish in phenol red-free DMEM (21063029, Gibco) containing 10% FBS. Twenty-four hours later, cells were washed twice in phenol red-free DMEM media and incubated in media containing D Red BSA (0.15 mg/mL) with or without FGF1 (5 nM) for 5 min at 37°C. Cells were placed on ice, washed twice with ice-cold media and then chased for 2 h in fresh media before fixation with 4% PFA. For dextran-TR experiments, cells were incubated in media containing dextran-TR (1 mg/mL) for 5 min at 37°C, washed 5 times with ice-cold PBS and fixed immediately, without a chase period, with 4% PFA. In some experiments, cells were treated with EIPA (75 µM) for 30 min at 37°C to block macropinocytosis.

Dextran fluorescence was quantified as described using ImageJ (National Institutes of Health)^38,39^. Briefly, maximum intensity projections of whole z-stacks were background subtracted using the rolling-ball algorithm. Images were then thresholded and segmented images of dextran-positive macropinosomes were measured using the “Analyze Particles” function with an area set to 0.2–20.0 μm^2^.

### Immunoprecipitation and immunoblotting

FRS2α−/− MEFs or FRS2α−/− PC12 cells and cells expressing the various mutant proteins were grown in 150-mm plates to confluency. Cells were left unstimulated or stimulated with FGF1 at the indicated concentrations and for the indicated times at 37°C. Following stimulation, cells were lysed as previously described^48^, and lysates were either analyzed directly by sodium dodecyl sulfate-polyacrylamide gel electrophoresis (SDS-PAGE) and immunoblotting or incubated with the appropriate antibodies overnight at 4°C. Immunocomplexes were resolved by SDS-PAGE and analyzed by immunoblotting with the indicated antibodies.

## Acknowledgments

This work was supported by 5P01AG051459-07 and RM1AWD0009897 grants from NIH and by Schlessinger’s laboratory discretionary funds.

