## Supplemental Figures for "Cellular responses to FGF1 are modulated by palmitoylation of the docking protein FRS2α"

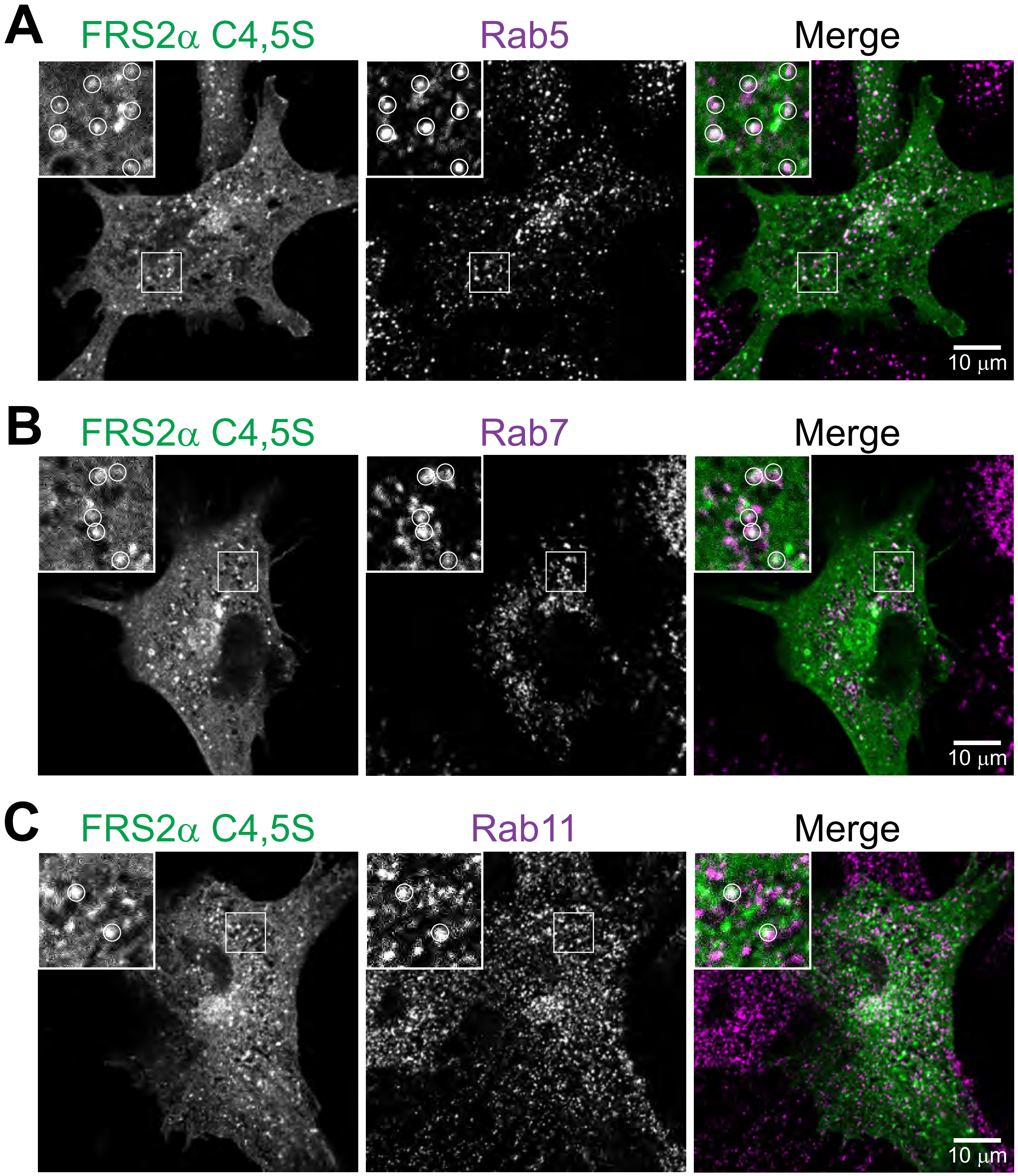


**Supplemental Figure 1. Colocalization of palmitoylation-deficient FRS2α C4,5S with endosomal markers.**

**(A–C)** Confocal images of FRS2α−/− MEFs transiently expressing FRS2α C4,5S-tdStayGold and immunostained for Rab5-positive early endosomes (A), Rab7-positive late endosomes (B) and Rab11-positive recycling endosomes (C). Individual confocal planes are shown. Insets show magnified views of boxed regions.

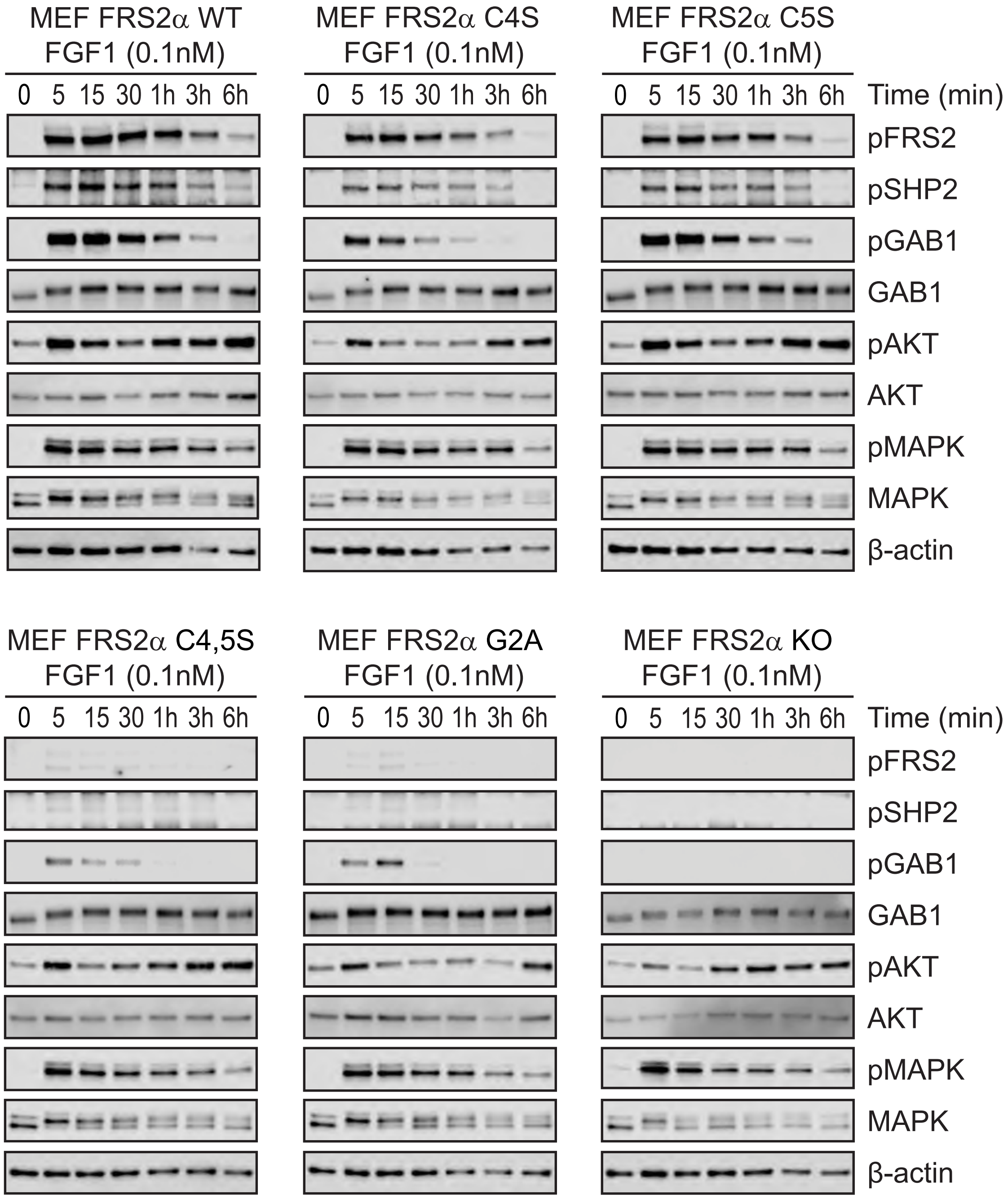


**Supplemental Figure 2. Kinetics of FGF1-induced signaling in FRS2α −/− MEFs and in MEFs rescued with FRS2α WT or its myristoylation and palmitoylation mutants.**

Unrescued FRS2α−/− MEFs and FRS2α−/− MEFs stably expressing FRS2α WT or its lipidation mutants were stimulated with FGF1 (0.1 nM) at 37 °C for the indicated times. Cell lysates were subjected to SDS-PAGE and analyzed by immunoblotting for tyrosine phosphorylation of FRS2α, SHP2, and GAB1, as well as for activation of MAPK and AKT using anti-pFRS2α, anti-pSHP2, anti-pGAB1, anti-pMAPK, and anti-pAKT antibodies, respectively. Immunoblotting with anti-GAB1, anti-AKT, anti-MAPK, and anti-β-actin antibodies was performed as a loading control.

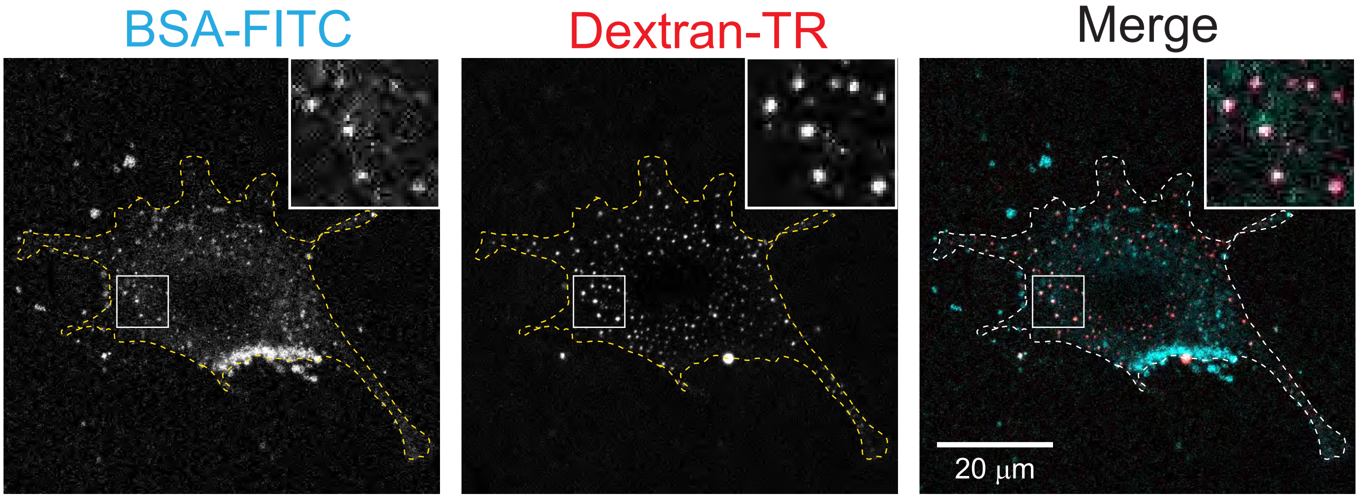


**Supplemental Figure 3. Macropinocytosis of BSA with dextran in MEFs.**

Confocal images of FRS2α−/− MEFs stably expressing FRS2α WT and stimulated with FGF1 (5 nM) for 5 min at 37 °C in media containing BSA-FITC and dextran-TR. Maximum intensity projections of z-stacks are displayed. Insets show magnified views of boxed regions.

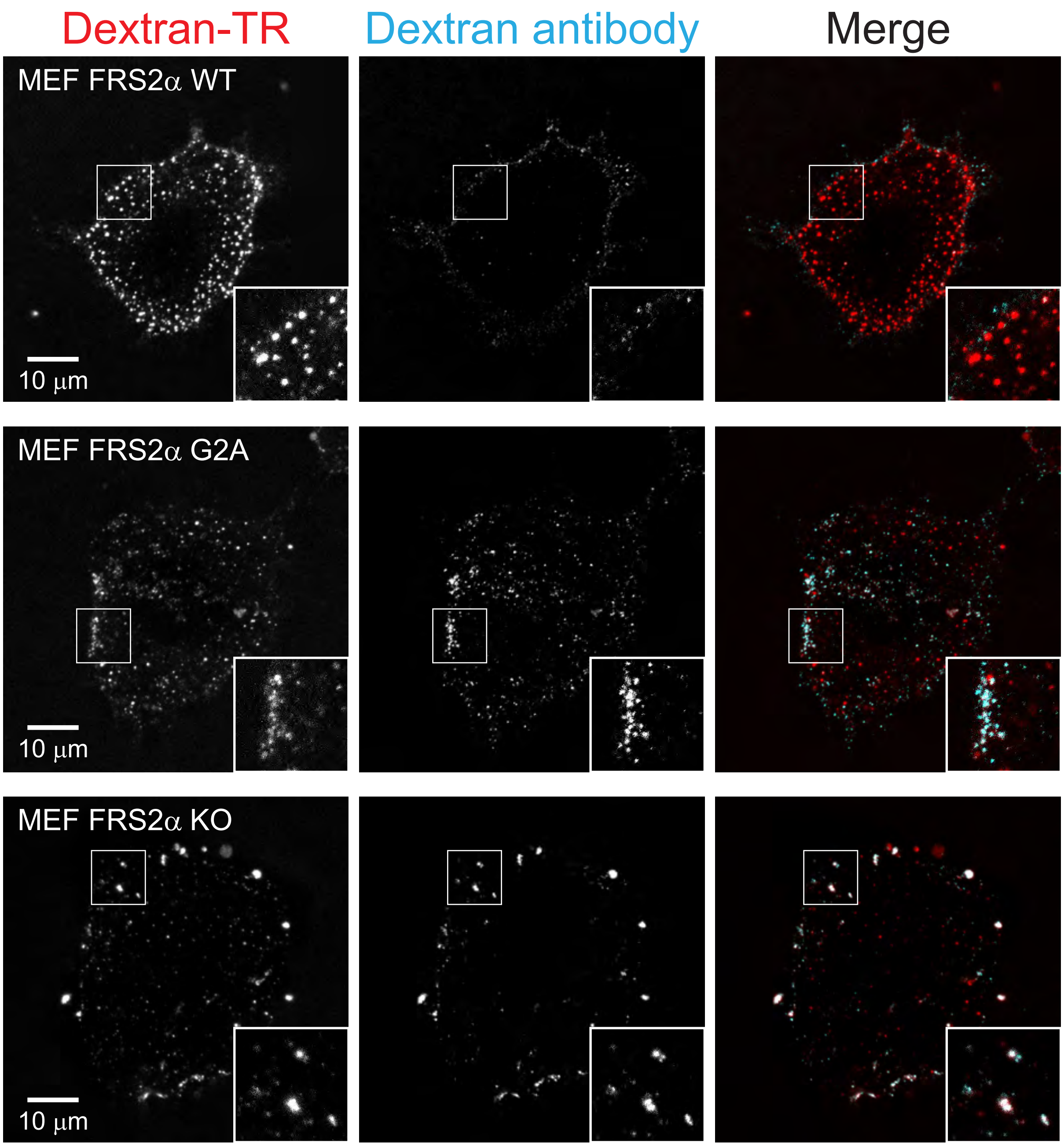


**Supplemental Figure 4. Macropinosome closure assay.**

FRS2α−/− MEFs left unrescued or stably expressing FRS2α WT or the palmitoylation deficient mutant FRS2α G2A were stimulated with FGF1 (5 nM) for 5 min at 37 °C, fixed (without permeabilization) and stained with anti-dextran antibodies and Alexa488-conjugated secondary antibodies. Cells were then imaged by confocal microscopy. Maximum intensity projections of z-stacks are displayed. Insets show magnified views of boxed regions.

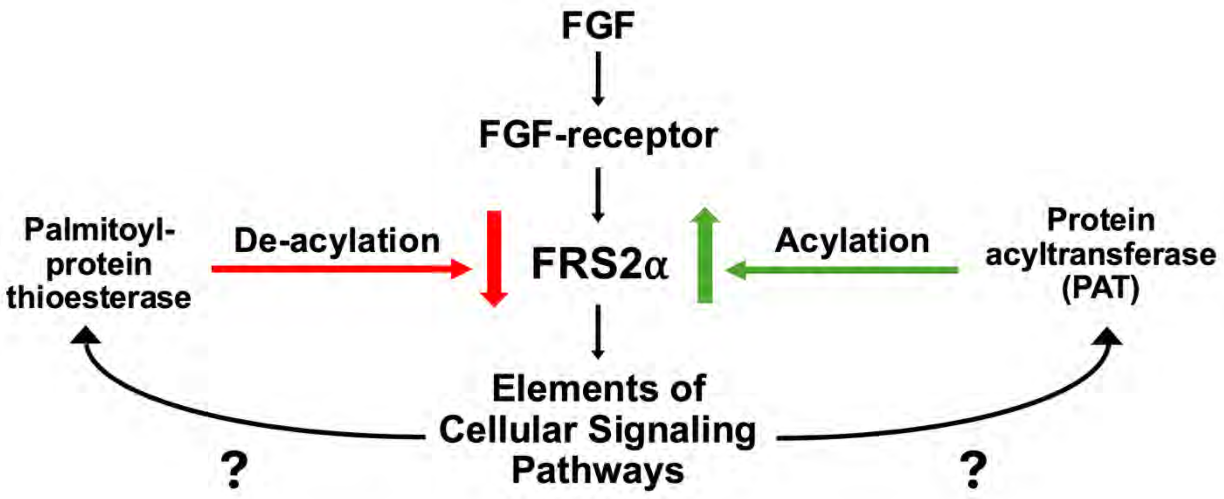


**Supplemental Figure 5. Putative mechanism for modulation of FRS2α palmitoylation by regulation of the activities of acylation and de-acylation enzymatic activities**.

Palmitoylation of FRS2α plays a critical role in regulating cellular signaling pathways stimulated by FGFs and neurotrophic factors. Accordingly, modulation of the activities of enzymes that acylate (protein acyltransferases) or de-acylate (palmitoyl protein thioesterases) FRS2α may represent an additional regulatory step controlling both the strength and spatial control of FRS2α-mediated signaling. We postulate that MAPK activation, or other downstream effectors stimulated by FGFs or neurotrophic factors, may phosphorylate these acylation and/or de-acylation enzymes, thereby creating additional positive or negative feedback loops that fine-tune ligand-induced cell signaling.
